# Viral host-range evolvability changes in response to fluctuating selection

**DOI:** 10.1101/771998

**Authors:** Morgan E. Mouchka, Dillon M. Dorsey, Genna L. Malcangio, Sarah J. Medina, Elizabeth C. Stuart, Justin R. Meyer

## Abstract

The concept of evolvability (the capacity of populations to evolve) has deep historical roots in evolutionary biology. Interest in the subject has been renewed recently by innovations in microbiology that permit direct tests of the causes of evolvability, and with the acknowledgement that evolvability of pathogens has important implications for human health. Here, we investigate how fluctuating selection on the virus, Bacteriophage λ, affects its evolvability. We imposed dynamic selection by altering the expression of two host outer membrane receptors. This, in turn, selected phage to alternately infect the host via a single, or multiple, receptors. Our selection regime resulted in two orthogonal evolutionary behaviors, namely enhanced or reduced evolvability. Strains with enhanced evolvability readily evolved between receptors, losing and gaining the ability to bind multiple receptors more quickly than the ancestral λ. This suggests the receptor-binding protein retained a genetic memory of past states and that evolutionary history can be used to predict future adaptation. Strains with reduced evolvability were refractory to re-specialization and remained generalists on both receptors. Consistent with this behavior, unevolvable strains had reduced rates of molecular evolution in the receptor-binding protein compared to their evolvable counterparts. We found a single mutation in the receptor-binding protein was sufficient to render these strains resistant to evolution and did so by counteracting a receptor-binding trade-off associated with generalism. In this way, cost-free generalization allowed for reduced evolution and evolvability while maximizing success in both environments. Our results suggest the response to fluctuating selection is contingent and can lead to distinct differences in evolvability. These findings contribute to a growing understanding of the causes and consequences of evolvability and have important implications for infectious disease management.

## Introduction

Evolvability is the capacity of populations to adapt through natural selection [1]. Because evolvability connects genotypes to adaptive variation, it is thought that evolvability itself is a quantifiable trait that is selected and maintained [2]. In this way, organisms vary not only in traits that determine their immediate fitness, but in their potential to generate better-adapted descendants. Because genetic variation drives adaptation, differences in evolvability stem from differences in the rate or effect of beneficial mutations between populations. Both theoretical and empirical work have demonstrated that genotypes with elevated mutation rates have greater evolvability under certain conditions [3–7]. For instance, multiple laboratory studies on *Escherichia coli* have found populations that evolve higher mutation rates experience increased rates of adaptation relative to populations lacking the mutator alleles [7, 8]. Research has also shown that evolutionary potential can vary based on differences in how mutations interact [2, 9–15] For instance, Woods *et al.,* [11] found that epistatic interactions between beneficial mutations caused differences in evolvability between subpopulations of *E. coli*, with these differences ultimately leading to the extinction of the less evolvable subpopulation.

Fluctuating environments are predicted to select for increased evolvability. In contrast to static environments where sustained directional selection results in optimization at local fitness peaks, the direction of selection in fluctuating environments is constantly changing, as are the location of the peaks. This puts pressure on populations to evolve characteristics that allow them to more efficiently explore new regions in the mutational landscape and keep up with the oscillating environment. While simulations have repeatedly demonstrated that changing environments select for more evolvable genomes [3, 4, 16–21], the majority of empirical studies measuring evolvability have done so in a stable, laboratory environment where selection for evolvability should be relatively weak. To fully understand the mechanisms by which fluctuating environments alter fitness landscapes and promote evolvability, evolvability should be measured empirically on organisms experiencing changing environments.

Understanding evolvability has important applications to processes ranging from protein engineering [22–24] to the rescue of populations threated by climate change [25–27]. One application for which evolvability is particularly relevant is predicting disease emergence. Emergence events are often the result of host range shifts whereby pathogens jump from their original host into a novel host species [28]. To make these shifts, pathogens must evolve to infect, adapt to, and be transmitted by the new host [29]. Viruses, owed to their large population sizes, short generation times, and high mutation rates, are particularly poised to respond to the selection pressures associate with host-range shifts [28, 30]. Indeed, multiple human pandemics including HIV from chimps [31] and the H1N1 ‘Spanish flu’ from birds [32] are the result of emerging viruses. During these shifts, viruses experience fluctuating environments as they coevolve with the new host and/or oscillate between hosts [33]. How do such fluctuating conditions affect viral evolvability? A better understanding of this question would enable effective surveillance strategies to anticipate emerging diseases.

Bacteriophage λ has long been a model for molecular genetics (Reviewed in [34]) and for the study of viral host-range evolution [35–39]. Experiments with λ allow for controlled selective environments, high replication, and evolutionary measurements over short periods of time, thus providing an ideal system to robustly measure evolvability. Bacteriophage λ normally infects *E. coli* via the host LamB receptor. Under conditions that promote lower expression of LamB, λ naturally evolves to exploit a novel receptor, OmpF. λ thus becomes a generalist phage that can utilize both receptors. It achieves generalization through mutations in its host-recognition protein, J. When the generalist λ is further evolved under conditions where cells only express one of the receptors, λ rapidly evolves enhanced function on the available receptor, and less function on the absent receptor [36]. This occurs because there is a trade-off between OmpF and LamB use and the shape of the relationship is such that there is a disproportionate cost for being a generalist [36].

Here, we build on this previous work by alternately selecting a generalist λ between conditions that favor receptor specialization on LamB and those that favor generalization on both receptors. A subset of replicate experiments yielded strains that were highly evolvable and readily switched back and forth between receptor specialization and generalization. The majority of replicates yielded strains that seemed refractory to selection for re-specialization, remained generalists, and were thus unevolvable. We sequenced the genomes of 48 evolved strains and found no specific molecular signature of the evolvable genomes. However, a single mutation in the receptor binding-protein, J, was sufficient to prevent strains from re-specializing. We also found that molecular evolution of the J protein was significantly decreased in unevolvable strains. We further investigated this mutation with the idea that understanding what prohibits evolvability may shed light on the contrapositive. We used genetic engineering to construct strains with specific genotypes to test hypotheses regarding how the mutation constrains evolvability.

## Results and Discussion

### Fluctuating environments select for two orthogonal strategies

We alternately evolved 12 replicate populations of the generalist ancestor between conditions selecting for receptor specialization and generalization (Figure 1A). Following this first round of selection (10 days), we isolated a representative λ from each population and selected those representatives back to generalization. At the completion of the second round, we isolated another representative from each population and continued this protocol for another full fluctuation. In the end, we isolated 48 unique λs that varied in the number of times they were pressured between receptor specialization and generalization.

**Fig. 1.**
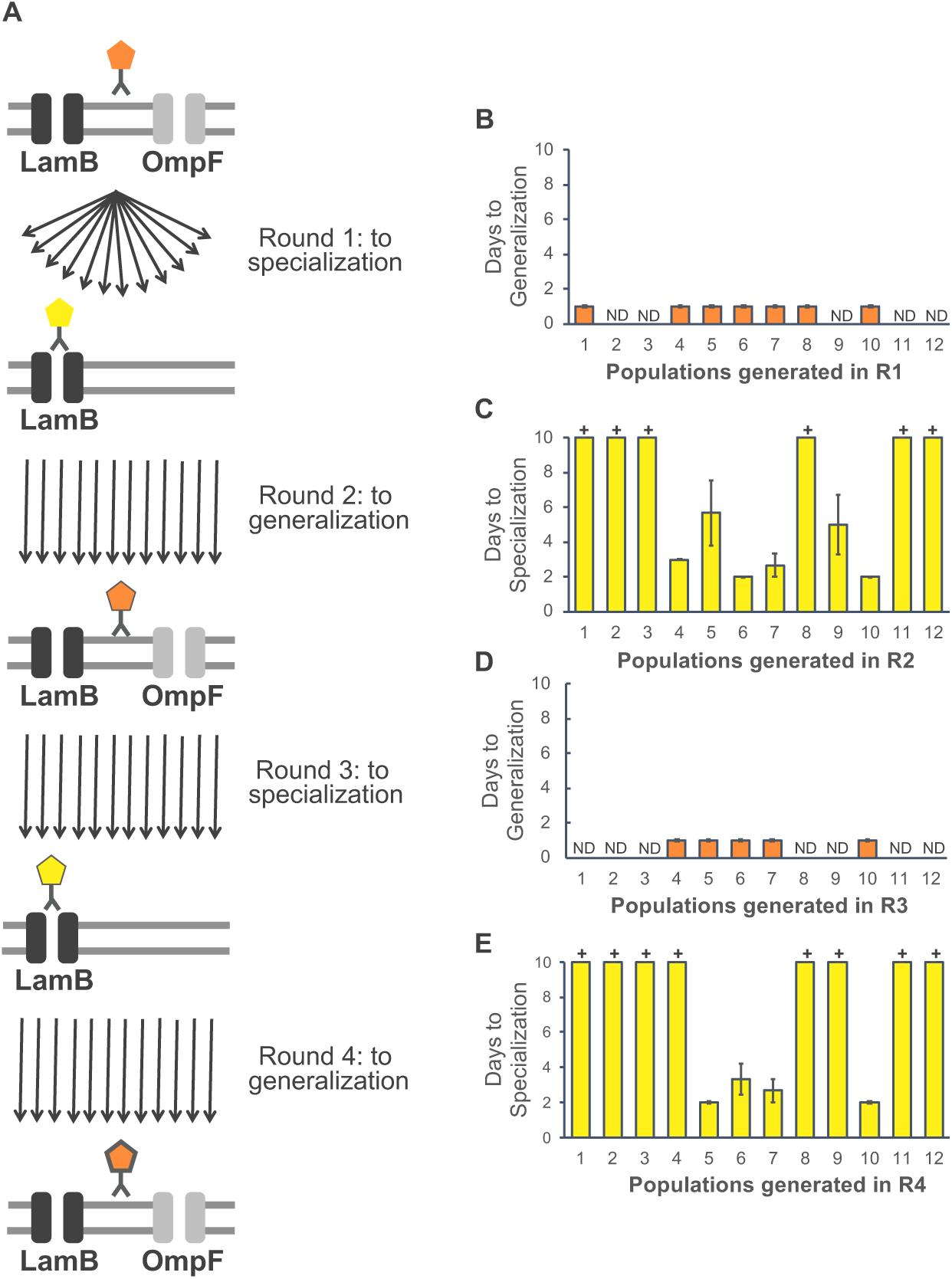
Generating and measuring evolvability. **(A)** To generate populations with varying degrees of evolvability, we subjected 12 independent populations of the generalist ancestor to successive rounds of evolution alternating between selection for receptor specialization (on the native receptor LamB) and generalization (on LamB and the novel receptor OmpF). At the end of each round of evolution, a single plaque was isolated from each population to seed the subsequent round. The 4 rounds yielded 48 strains, one per round from each population, with 24 strains evolving toward specialization and 24 evolving towards generalization. **(B-E)** Evolvability (number of days to evolve specialization or generalization) of the strains generated in Rounds 1-4 (R1-R4) (n = 3, average ± standard error). Several generalist strains did not evolve back to LamB specialization within the 10-day time frame of the replay experiment (+ on graphs in Fig. 1D and F). Because these strains remained generalists, they could not be assayed in subsequent replay experiments (ND on graphs in Fig. 1C and E).

To determine if and how successive rounds of fluctuating evolution altered evolvability, we subjected the strains to generalizing (Rounds 2 and 4) or specializing (Rounds 1 and 3) conditions and measured the number of days it took them to evolve back to generalization (Figure 1C and 1E) or specialization, respectively (Figure 1D and 1F). Two distinct phenotypes emerged; those that responded to selection and readily evolved back and forth between receptor specialization and generalization (hereon referred to as evolvable strains), and those that were resistant to selection and remained ‘stuck’ as generalists (hereon referred to as stuck strains).

The evolvable strains were, indeed, highly evolvable, taking a single day to regain OmpF function, regardless of population or round (Figures 1C and 1E). Note that several strains (ND in Figures 1C and 1E) could not be assayed for evolvability towards generalization because they remined generalists during previous rounds of evolution. In previous work exploring the evolution of novel receptor usage, the ability of λ to gain OmpF function *de novo* took 9-17 days, a median of 12 days, and occurred in only 25% of the populations [35]. Comparatively, our results demonstrate restoring OmpF function occurs quickly (within 1 day) and is highly repeatable (occurred in 100% of assayed populations).

Evolution toward LamB specialization took longer or did not happen at all (Figures 1D and 1F). For the evolvable strains, there was significant genetic variation among the isolates in how long they took to re-evolve specialization (ANOVA F = 3.286, p < 0.0071), and the overall pattern is that the more evolved strains took less time (Figure S1; Linear regression F = 5.6212, p < 0.0418), suggesting that the lineages gained evolvability through the fluctuations.

The dominant pattern observed was that half of strains generated in Round 2 and two-thirds of strains generated in Round 4 did not evolve back to specialization. These findings were unexpected for two reasons. First, we hypothesized that fluctuating conditions would select for more evolvable strains. Instead, increasing rounds of evolution led to a larger number of strains that were resistant to selection. Second, we expected that re-gaining function would be more difficult than losing function, however; regaining OmpF use happens more quickly than losing it and some strains are completely resistant to losing this function.

The two orthogonal evolutionary behaviors in replicates that are otherwise identical reveals the idiosyncratic nature of evolution. What drives these alternative behaviors? One hypothesis is that λ evolution is historically contingent; such that early mutations and adaptations affect later evolutionary trajectories [40].

### Genetic signatures of evolvability and evolutionary resistance

To test the historical contingency hypothesis, we sequenced the genomes of the 48 strains to identify mutations that dictate λ’s evolutionary path. We identified 41 unique mutations across the 48 strains, with 71% of unique mutations occurring in coding regions. The majority of coding mutations (82%) occurred in the reactive region of λ’s host-recognition protein, J. As this was the most informative region of the genome, we narrowed our focus to the J gene. All but one of the mutations in J were non-synonymous, implying positive selection for beneficial changes.

Figure 2 shows the mutation matrix for the J gene arranged by evolvable or stuck populations/strains. For evolvable populations, there is no single mutation or set of mutations that defines evolvability. Instead, multiple and diverse mutations enable these strains to readily evolve between specialization and generalization. These mutations often evolve in parallel, with a single mutation or set of mutations arising independently in multiple populations (*e.g.,* g3152t in populations 1, 7, and 10, g3226a in populations 5, 6, and 10, a3319g in populations 4 and 6, and t3364c, t3320c and t3380c in populations 7 and 10). The fact that independent lineages evolved the same mutation provides strong support for the selective advantage conferred by these mutations under a particular selection regime. Indeed, parallel mutations often arise during the same round of evolution, suggesting they are specific to the direction of selection. For instance, a3119g evolves in both populations 4 and 6 when the strains are being evolved toward specialization on LamB (Round 1).

**Fig. 2.**
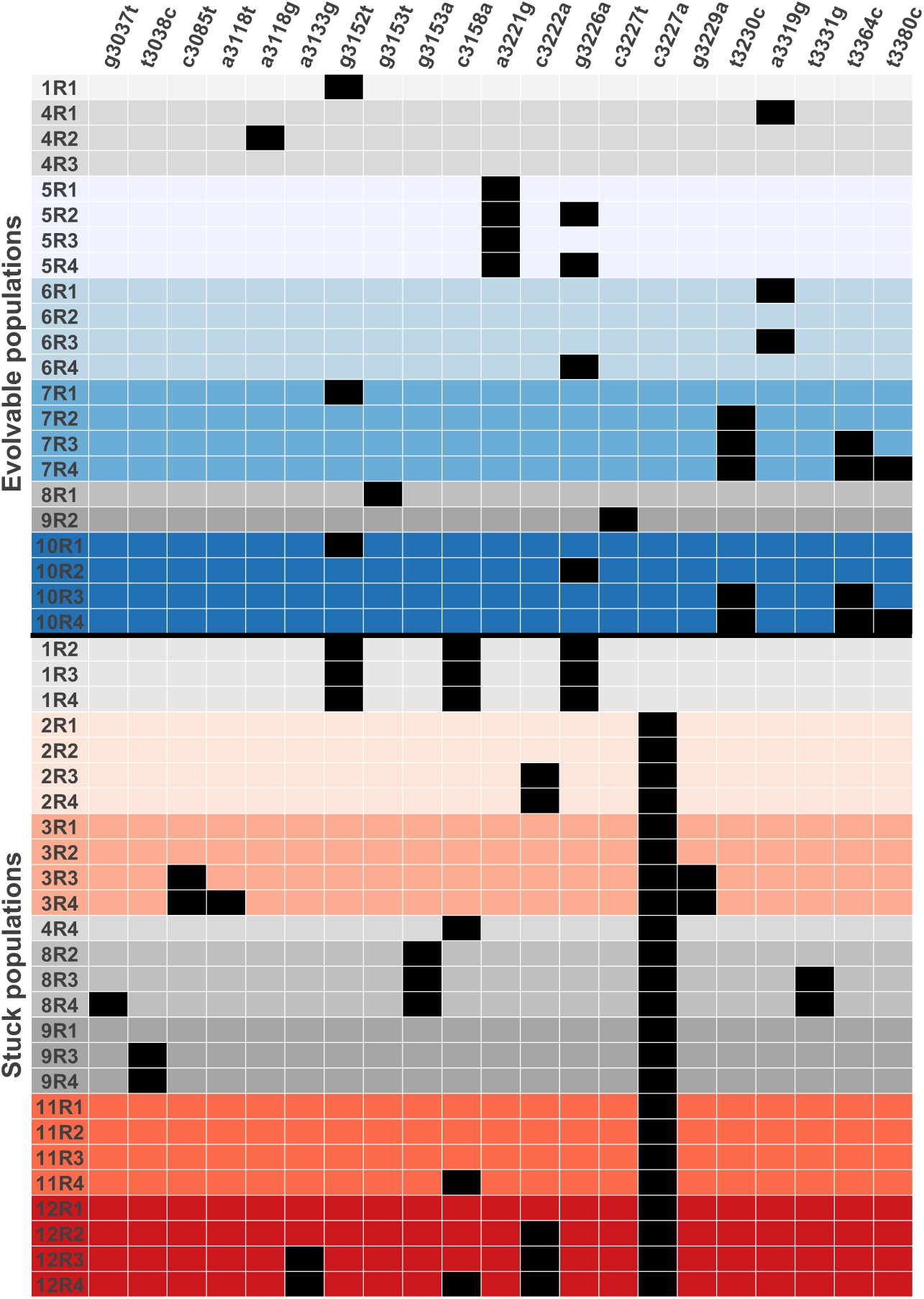
Genetic signatures underlying evolvability. Substitutions in the *J* gene compared to the ancestor from λ populations subjected to 4 rounds of alternating evolution between receptor specialization (LamB) and generalization (LamB and OmpF). Populations/round are rows; position/mutation are columns. Populations that readily evolved between specialization and generalization are represented by shades of blue, with each shade denoting an independent population. Populations that became ‘stuck’ as generalists after the first round of evolution and persisted in that state for subsequent rounds are represented by shades of red. Populations that alternated between ‘stuck’ and evolvable states are represented in gray. Except for one population (population 1), populations that become ‘stuck’ as generalists have a cytosine to adenine substitution at position 3227.

The presence of 8 reversions at 4 positions across 5 populations demonstrate that reversions are a common feature of the evolvable strains. In contrast, the stuck strains exhibit only a single reversion (t3227a in population 9). In some instances, double (g3226a in population 5) and even triple reversions (a3319g in population 6) occur when a mutation is gained and lost multiple times over successive rounds of alternating selection. Similarly, Crill et al [41] found that alternating *ɸ*X174 phage between an ancestral and novel host resulted in reversions in 11 of the 55 sites that underwent substitution, including multiple reversions at a single site upon host shifts.

As second-site compensatory mutations are proposed to arise more often than reversions [42], there are two possible explanations as to why reversions commonly arose in our experiment. First, large population sizes have been shown to facilitate the evolution of reversions [42], and this is especially true under changing selection regimes (Schrag et al 1997 [43]. Although the densities of our phage varied by strain following 24 hours of evolution, populations were large and had greater than 1 x10^9^ plaque forming units (data not shown). Second, the use of reversion to evolve between selection regimes suggest these particular mutations lead to antagonistic pleiotropy between receptor generalization and specialization. This is consistent with the idea that there are trade-offs associated with receptor tropism [33, 36, 41], such that mutations that are beneficial in one environment limit fitness in an alternative environment. Interestingly, only one of the 4 positions where reversions occur is a conical mutation associated with the evolution of novel OmpF function [44]. This implies that evolvable strains can use both previously explored and novel mutational pathways to gain and loose receptor function.

Unlike for the evolvable strains, a single mutation or set of mutations conferred resistance to selection for stuck strains (Figure 2). For population 1, a set of up to 3 mutations (g5152t, c3158a, g3226a) prevented strains from evolving back to specialization. Since strains with just one of these mutations remain evolvable, yet strains with all three mutations are stuck, we can infer at least two mutations are necessary to prevent evolution back to specialization. However, lacking strains exhibiting varying combinations of these three mutations, we are unable to determine if all three mutations are necessary and/or sufficient to prevent re-specialization.

For all the other 8 populations where strains became stuck, a single cytosine to adenine substitution at position 3227 conferred resistance to evolution. Indeed, as there are several stuck strains (2R1, 3R1-3R2, 11R1-11R3, 12R1) that differ from the ancestor by only this single mutation, we can conclude that this mutation is sufficient to render these strains stuck. The parallel evolution of this mutation in multiple independent lineages suggests that it is highly adaptive under fluctuating conditions. Interestingly, the only reversion in a stuck strain occurred at this site in population 9, and its loss and regain mirrored its phenotypic behavior (Figure 1C and D). That is, losing the mutation led to evolvability, while regaining the mutation led to the inability to re-specialize. This provides further evidence that this mutation is sufficient to cause the stuck phenotype and that it confers evolutionary resistance.

Given the distinct phenotype and genotypes of stuck strains, we next wanted to determine if these strains were also ‘stuck’ at the molecular level. To do this, we compared the average number of J mutations per round between evolvable and stuck strains (Figure 3). We found the J mutation rate of stuck strains was 47% lower than that of evolvable strains, suggesting a dampened rate of molecular evolution in strains stuck at generalists. As such a decrease in substitution rate could have major implications for evolution over evolutionary timescales, we next turned our attention to determining why the stuck phenotype arises. To do this, we focused on the c3227a mutant so that the effect of the single mutation could be isolated.

**Fig. 3.**
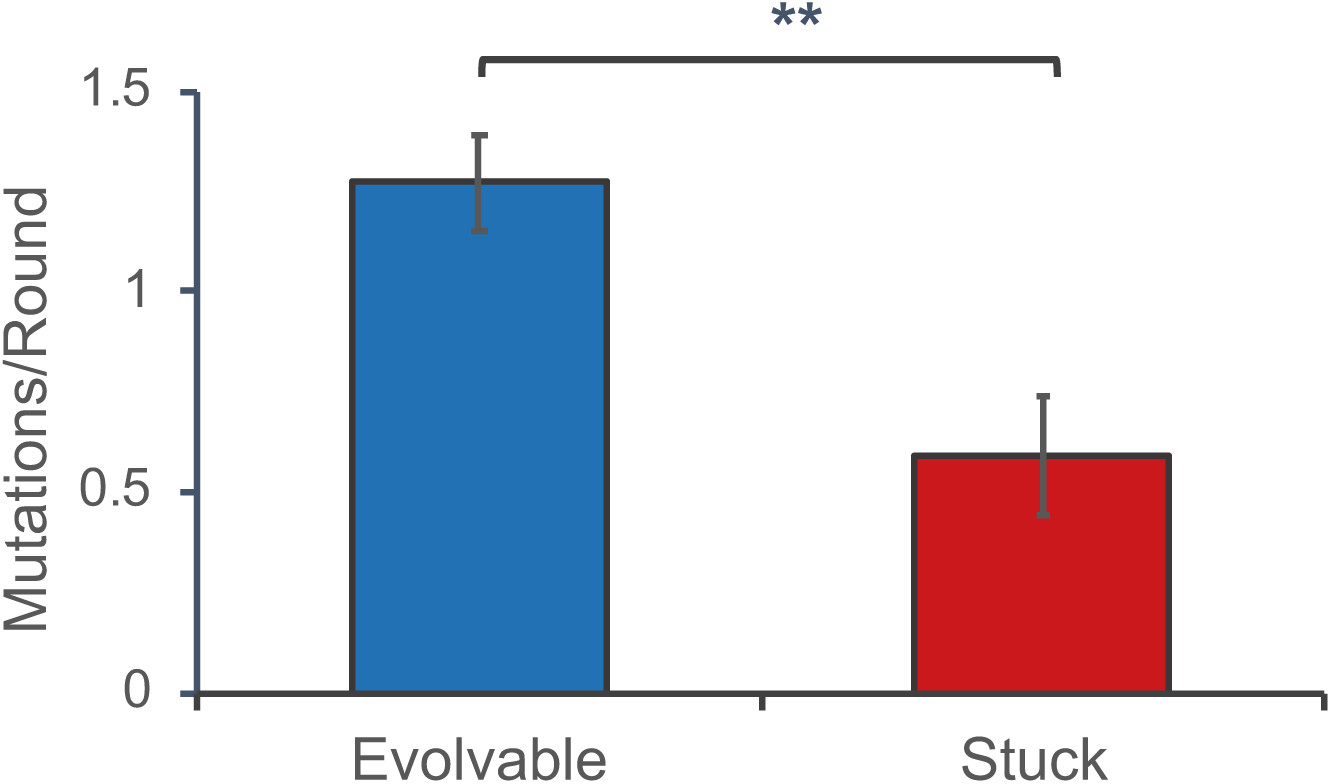
Number of J mutations per round (average ± standard error) for stuck and evolvable strains. (Kolmogorov-Smirnov test: D = 0.578, with Monte Carlo permutation p < 0.001).

### Epistasis and trade-offs

The majority of strains experiencing fluctuating conditions become resistant to selection and remain generalists. Given the trade-off between OmpF function and reduced binding on LamB, what could explain the evolution and persistence of these stuck strains? We developed three hypotheses to explain the stuck genotype/phenotype: 1) Negative epistasis between the stuck and evolvable mutations prevents the evolvable mutations from conferring a fitness benefit, thereby their emergence, 2) Masking epistasis by the stuck mutation that blocks the effect of evolvable mutations, and 3) the stuck mutation breaks the receptor-binding trade-off to generate a genotype that is adept at using both receptors and is thus refractory to selective pressures.

#### Negative epistasis

Epistasis has been shown to play an important role in viral evolution, in part because viral proteins are often highly interactive and multifunctional [13, 45, 46]. Indeed, both theoretical and experimental research has demonstrated that epistasis can strongly constrain evolvability [10-13, 15, 47, 48]. Negative epistasis occurs when two mutations lead to a less fit phenotype than the two individual mutations alone confer. Negative epistasis between the stuck mutation and those associated with evolvability could prevent stuck lineages from re-specializing on LamB. This, in turn, could limit evolutionary trajectories for the stuck linages and lead to reduced rates of molecular evolution. Looking at the mutation matrix (Figure 2), there is very little overlap between mutations that arose in evolvable and stuck strains. In fact, there is not a single mutation that occurs in both evolvable strains and strains with the canonical stuck mutation. This pattern suggests negative epistasis between stuck and evolvable mutations such that the two states are incompatible.

To test this hypothesis, we used multiplexed automated genome engineering (MAGE) [49, 50] to edit a lysogenic phage and construct alleles corresponding to the ancestral generalist strain, a strain with the conical stuck mutation (c3227a), a strain with a mutation linked to evolvability (a3319g), and a strain with both the stuck and evolvable mutations. The ‘evolvable’ mutation a3319g was chosen because it arose multiple times in different lineages, it was the only mutation present in these strains meaning it must have caused the specialization, it only evolved during selection for specialization, and it only occurred in strains that were evolvable. We then took each of the constructed strains and competed them against a genetically marked ancestral strain to measure relative fitness. We hypothesized negative epistasis between the stuck and evolvable mutation would lead to phage that was either inviable (*e.g.,* incapable of forming a functional particle) or had very low competitive fitness. Figure 4 shows the competitive fitness of the constructed strains relative to the genetically marked ancestral strain. Contrary to our expectations, phage populations with both mutations were not only viable, but highly fit. After normalizing for the fitness of the ancestral strain, there was no significant difference in fitness between the construct with both mutations, and the additive fitness of the constructs with either the stuck or evolvable mutation (Wilcoxon Rank Sum, Z = 1.1619, p < 0.2543). Thus, we concluded that the mutations are not genetically incompatible and that negative epistasis between them does not explain why stuck strains are constrained in their evolvability.

**Fig. 4.**
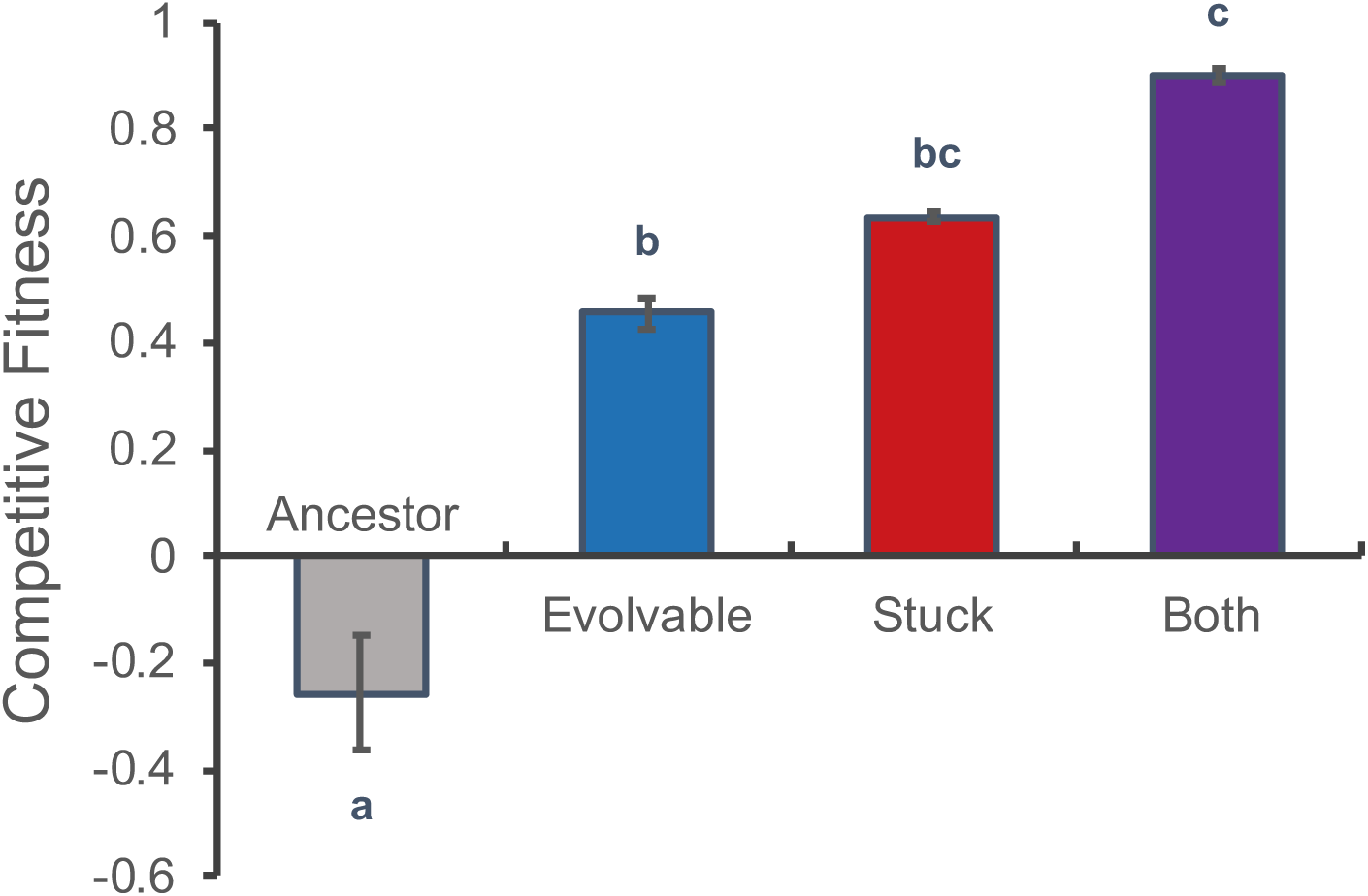
Competitive fitness (n=2) of phage engineered with an ‘evolvable’ mutation that arose in multiple strains, the canonical ‘stuck’ mutation, both the ‘evolvable’ and ‘stuck’ mutations, and an ancestral control (average ± standard error). Strains were competed against a genetically marked ancestral strain. (ANOVA: F-ratio = 76.67, p < 0.0005. Differing letters denote significance via a Tukey-Kramer HSD test. See Table S2 for pairwise p-values).

#### Masking epistasis

The inability of stuck strains to re-specialize on LamB could be limited by epistasis that masks the phenotypic effects of the evolvable mutations by the stuck mutation. To test this hypothesis, we quantified the effect the evolvable mutation has on λ’s receptor specificity when it is engineered into an ancestral genome versus a genome that has the stuck mutation. The prediction is that the evolvable mutation’s effect will be blocked, or at least diminished in the second genome. To quantify λ’s receptor specificity, we measured the relative ability of phage to use each receptor and calculated a specialization index that ranges from 0 (perfectly symmetric generalist) to 1 (complete specialist on LamB) [36]. Figure 5 shows the specialization on LamB for the constructed strains. Contrary to our hypothesis, we found the construct with both mutations was nearly a complete specialist, much like the evolvable construct. This implies, if anything, that the evolvable mutation masks the phenotypic effects of the stuck mutation, functionally enabling a strain with the stuck mutation to regain evolvability. Masking epistasis by the stuck mutation therefore does not explain the inability of stuck strains to regain specialization on LamB.

**Fig. 5.**
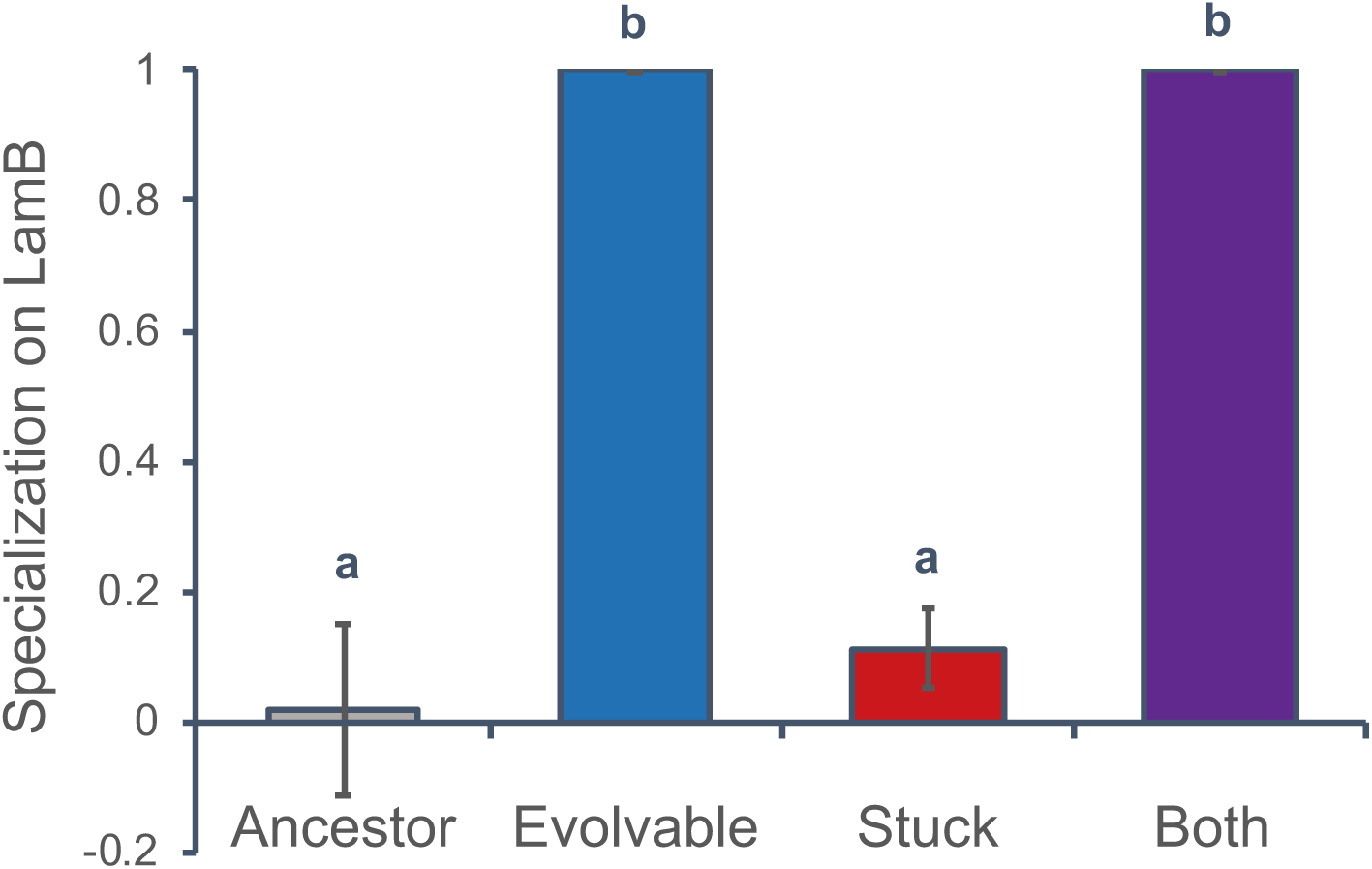
Specialization index (n=3) on LamB for the engineered phage strains (average ± standard error). The index is a measurement of the relative ability of a phage to infect via a single receptor. Index of values of 1 and 0 indicated complete specialization on LamB and compete generalization, respectively. (Kruskal-Wallis Rank Sum: *χ*^2^ = 17.45, p < 0.0006. Differing letters denote significance via a Steel-Dwass test. See Table S3 for pairwise p-values).

#### Breaking the trade-off

Host-range expansion in viruses often results in reduced fitness in the ancestral host, implying that generalism comes at a cost [41, 51, 52]. To explain why strains would resist specialization given the potential cost of generalization, our third hypothesis posits that the stuck mutation enables strains to break this receptor-binding trade-off. In this way, the stuck mutation results in a superior generalist that can maintain OmpF function while more effectively binding LamB. In a highly fluctuating environment, strains with the stuck mutation remain generalist at little to no cost and no longer need to respond to shifting selection pressures.

The first indication that the stuck mutation may provide enhanced use of both receptors is that it arose when populations were being selected towards both specialization and generalization. Although, this observation may also indicate that the mutations improve some other generic quality of J. To test whether it enhances receptor use, we measured receptor adsorbtion rates of the ancestor and the strain with the stuck mutation. We also measured adsorbtion rates for the strain with the evolvable mutation as an additional point of comparison. The stuck strain breaks the trade-off and increases binding on both LamB and OmpF compared to the ancestor (Figure 6). In contrast, the evolvable mutation improved adsorbtion to LamB and reduced adsorbtion to OmpF. The stuck mutation does not improve adsorbtion on LamB as much as the evolvable mutation, however, the additional gain by the evolvable mutation seems unnecessary for λ’s evolution towards specialization. A re-evaluation of Figure 4 shows that there is no fitness difference between genotypes that have the stuck or evolvable mutation when they are challenged to re-specialize (Wilcoxon Rank Sum, Z = −1.1619, p < 0.2453). An interesting consequence of breaking the trade-off is the rate of molecular evolution was lower for stuck strains (Figure 3). If the stuck mutation results in a superior generalist that no longer needs to alter phenotypes to maximize function in both environments, there is less need to respond to selection and evolve additional mutations.

**Fig. 6.**
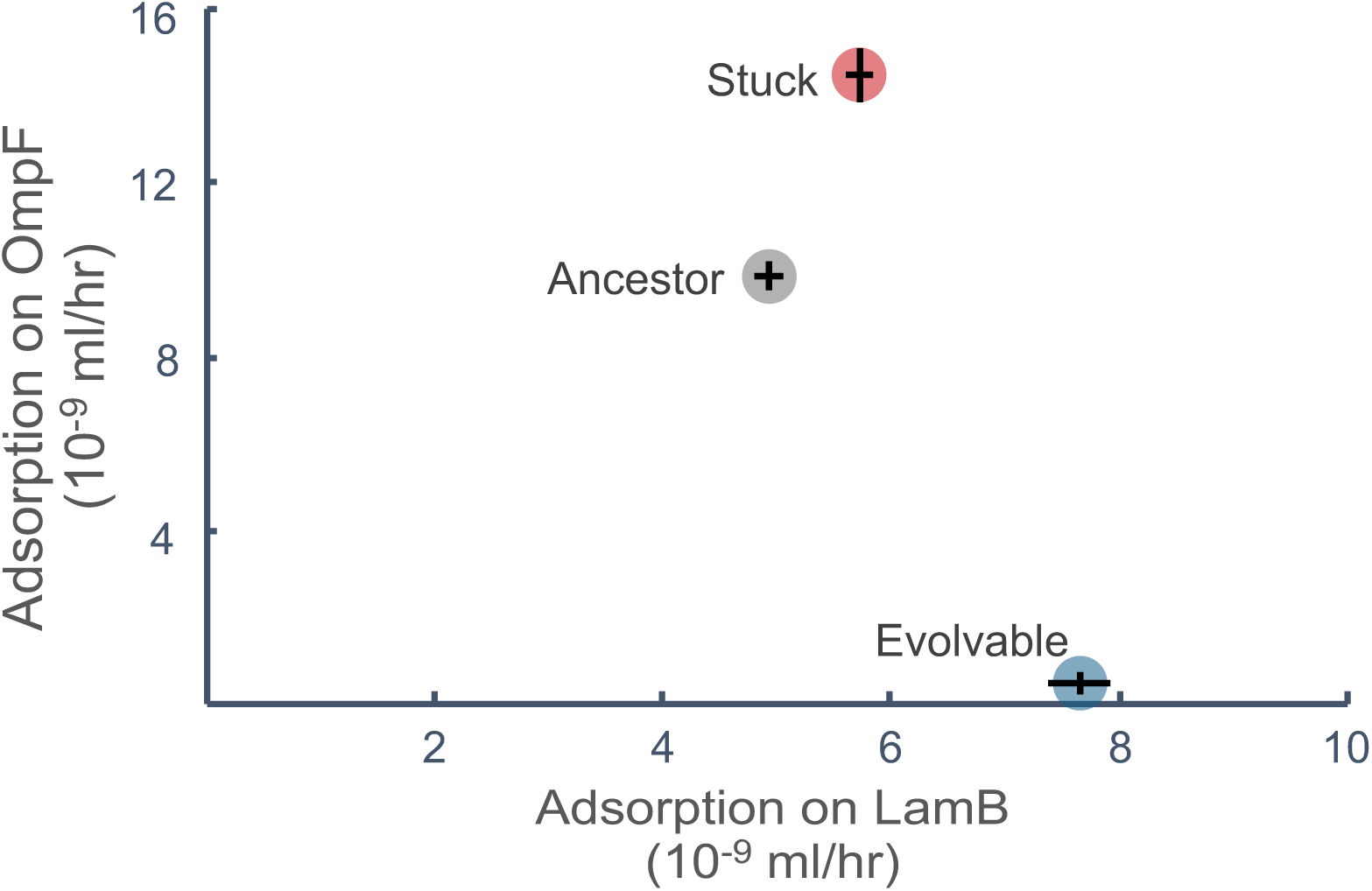
Adsorption rates (n = 9) on LamB and OmpF for engineered phage strains (average ± standard error). Adsorption rates on different receptors are plotted on the same graph to show relationships, but were collected in independent trials. (Kruskal-Wallis Rank Sum for LamB: *X*2 = 22.59, p < 0.001 and OmpF: *X*2 = 23.15, p < 0.001. Pairwise comparisons between constructed strains were all highly significantly different for both LamB and OmpF adsorption rates using a Steel-Dwass test. (See Table S4 for pairwise p-values).

While trade-offs are thought to be ubiquitous in evolutionary biology, they are not necessarily constitutional. Under certain circumstances, trade-offs are caused by a biophysical constraint that cannot be broken without disrupting physical laws. However, the majority of trade-offs are likely caused by evolutionary constraints that may be broken if enough genetic variability can be generated to counteract costs. Indeed, numerous studies have found that while organisms incur a cost associated with the evolution of a novel trait, this cost can be diminished or eliminated by further evolutionary change [43, 53–57]. For instance, McKenzie et al., [54] found that Australian sheep blowflies with a mutation conferring insecticide resistance developed more slowly and had reduced survival compared to their sensitive counterparts in the absence of insecticide. However, this cost of resistance was soon eliminated by a second mutation that made the resistant flies as fit as their sensitive counterparts. It appears that the stuck mutation acts similarly, breaking the receptor-binding trade-off and diminishing, if not eliminating the cost of generalization.

### Evolvability and genetic memory

Evolvable strains repeatedly lost, and subsequently regained, receptor function and did so more readily with increasing numbers of fluctuations. What is the mechanism underlying this increased evolvability? Do these strains exhibit a generic enhancement of evolutionary potential? Or do they retain an evolutionary ‘memory’ allowing adaptations to previous conditions to be remembered and recaptured more quickly [58]? To address these questions, we tested whether the generalists had simply re-gained OmpF function or had also evolved to use an alternative receptor. *E. coli’s* outer membrane has several proteins with similar structures to LamB and OmpF, and a closely related lambdoid phage infects through one of these proteins (OmpC) [59]. Evolution to use a third receptor is therefore possible and could be driven by generic evolvability. However, we found that none of the generalist strains tested could infect *E. coli* via an alternative receptor, supporting the evolutionary memory hypothesis.

In developmental biology, evolutionary memory is thought to span lineages and time, enabling re-expression of developmental pathways to regain function. For instance, Whiting *et al.,* [60] showed that stick insects (order Phasmatodea) diversified as wingless insects and that wings were derived secondarily on multiple occasions via the re-expression of a developmental pathway that evolved only once and early during insect diversification. Over shorter evolutionary timescales, retaining a memory of past states results in more evolvable populations. For instance, recent work in digital organisms shows that fluctuating conditions alter the genetic architecture of populations, making them better able to adapt to previously seen conditions [18].

While it is widely accepted that memory could be stored in developmental pathways and gene regulatory networks through the reactivation of latent genes, how might evolutionary memory be stored at the protein level? Accumulating mutations can change the scaffolding of proteins to enable novel functions. Natural selection can then act on this existing scaffold to more readily alternate between functional states. In this way, a past state can more easily be ‘remembered’ and accessed with fewer mutations, without rebuilding the scaffold from the ground up. This memory of past functions has important implications for viral host range shifts and reemergence. Viral strains that carry vestigial information from previous host environments are more likely to successfully switch between hosts. By superimposing host switches onto phylogenetic reconstructions of viral strains, we can use past evolutionary states to predict which strains are more likely to reemerge.

### Conclusions

We found fluctuating *λ* between receptor specialization and generalization resulted in two distinct strategies that either enhanced or reduced evolvability. Strains with enhanced evolvability readily and repeatedly evolved between specialization and generalization and gained evolvability with increased numbers of fluctuations. This implies the receptor-binding protein retained a memory of past evolutionary states, and that much like for gene regulatory systems, genetic memory can be stored in proteins. This has important implications for emerging disease surveillance. If past evolutionary states can predict future outcomes, phylogenic reconstructions of viral host histories could be used to predict the evolvability of extant strains and identify those most likely to reemerge.

The outcome of fluctuating conditions is contingent, however, and the same conditions that promote evolvability can also favor traits that reduce evolvability while enhancing within-generation performance. Indeed, the majority of strains were refractory to re-specialization and evolved adaptations that facilitated superior generalism by counteracting the receptor-binding trade-off. This cost-free generalization, in turn, reduced evolvability and slowed evolution. The long-term evolutionary implications of this strategy are unclear. On one hand, the evolution of generalism is thought to foster improved performance in novel environments [61] and flexible receptor-binding proteins could facilitate further host-range shifts. On the other hand, the reduced standing genetic diversity in these refractory strains could hinder future adaptation.

While much work is to be done on this subject, our research contributes to a growing understanding of the causes and consequences of evolvability. We provide novel and empirical evidence for how genomic features can impact the evolutionary capacity of organisms and how past evolution can impact future adaptation. Our results also have practical implications relevant to translational research. Viral evolvability is something to be both countered, as in the case of antiviral drug development, or exploited, as in the case of phage therapy. Our results suggest that evolvability can be reduced or enhanced under identical conditions and that viral genomes could be manipulated to constrain or augment further evolution. Lastly, our results are applicable to the study of infectious disease and have the potential to inform predictive frameworks for viral host-range shifts and disease emergence.

## Materials and Methods

### Media

Media were prepared according to the following specifications. **LB** (Lennox Broth): 10 g tryptone, 5 g yeast extract, and 5 g sodium chloride per liter water. **TrisDM:** 0.28 g dipotassium phosphate, 0.08 g monopotassium phosphate, 1 g ammonium sulfate per liter water, supplemented with a final concentration of 50 mM Tris (pH 7.4), 6 *μ*M thiamine, 0.02% LB, 0.2 mM calcium chloride, 20 mM magnesium sulfate, and either 6 mM maltose or glucose. **LBM9**: 20 g tryptone and 10 g yeast extract per liter water, supplemented with a final concentration of 47.7 mM disodium phosphate, 22.0 mM potassium phosphate monobasic, 18.7 mM ammonium chloride, 8.6 mM NaCl, 0.2 mM calcium chloride and 20 mM magnesium sulfate. **LB agar**: 10 g tryptone, 5 g yeast extract, 5 g sodium chloride, and 16 g agar per liter water. **Soft agar**: 10 g tryptone, 1 g yeast extract, 8 g sodium chloride, 7 g agar, 0.1 g glucose per liter water, supplemented with a final concentration of 2 mM calcium chloride and 10mM magnesium sulfate. **Deoxy-galactose agar**: 5.34 g potassium phosphate dibasic, 2 g potassium phosphate monobasic, 1 g ammonium sulphate, 0.57 g sodium citrate, 16 g agar, 0.12 g magnesium sulphate, 0.0002% w/v thiamine, 0.0004% w/v biotin, 16 ml glycerol, and 0.2 g 2-deoxygalactose per liter water.

### E. coli strains and culturing

Several *E. coli* K-12 derivatives were used in this study. These include strains from the Keio knockout gene collection (OmpF^—^ JW0912, LamB^—^ JW3996, and the undisrupted parental *wildtype* strain BW25113) [62] that were used for culturing *λ*, infused overlay and spot plates, evolution experiments, and phenotypic assays. To determine if λ had gained function on additional receptors, we plated on an OmpF^—^LamB^—^ double knockout that we engineered (see *Genetic Engineering* below). DH5*α* was used for plasmid cloning (Subcloning Efficiency™ DH5*α*™ Competent Cells; Invitrogen) and for plating the competitive fitness assays. We used HWEC106 (provided by Harris Wang, Columbia University) to genetically engineering λ. HWEC106 has a *mutS* deletion by insertion of the *bla* beta-lactamase resistance gene and carries the λ-red recombineering plasmid pKD46 [63]. Lastly, we used SYP042, which contains a plasmid encoding a phage recombination (R) gene fused to lacZ*α* [38, 64, 65] to construct a phage with a visual marker when grown on lacZ*α*^—^ host cells.

*E. coli* strains were stored frozen at −80°C with 15% v/v glycerol. Cultures were grown from ∼2 *μ*L of frozen stock in sterile glass tubes with 4 mL of LB media (unless indicated otherwise below) and at either 30°C (HWEC106) or 37°C (all others) at 220 rpm overnight. When culturing HWEC106, SYP042, or DH5*α* with plasmid, 50 *μ*g/mL ampicillin (Fisher Scientific) was added to the media.

### *λ* strains and culturing

λ has two life cycles; the lytic phase, where infection leads to rapid phage replication and cell death, or lysogeny, the stable incorporation of the λ DNA into the host genome [66]. The base strains used in our study have particular mutations in the *cI* gene that control the lysogeny-lysis switch. The ancestral *λ* is derived from the strictly lytic base strain, cI26 [67], and had previously evolved function on the novel OmpF receptor [35]. The ancestral *λ* also has a unique region in the J gene resulting from a previous recombination event with an *E. coli* host. The recombination is 1336 bp long from 1545 to 2796 in J, and includes an extra 84 bp. The ancestral *λ* strain was used for the fluctuating evolution experiments. cI857 (provided by Ing-Nang Wang, State University of New York at Albany) is a base strain that can alternate between the lysogenic and lytic life-cycles due to temperature-sensitive mutation in the cI protein. If the temperature is low, *λ* remains integrated in the *E. coli* genome. When heat shocked, integrated cI857 lysogens are induced and lyse their host, releasing infectious *λ* particles. We used this strain for genetic engineering because the lysogen is easier to genetically modify [36]. Furthermore, lysates induced from lysogens only takes ∼2 hrs, (unlike a typical overnight co-culture with host cells that can take 16 hours), creating newly induced phage at similar stages in the lifecycle. See Table S6 for list of *λ* strains.

Lysates from lysogens were produced by inoculating 133 *μ*l of a lysogen culture into 4 mL of LBM9 and growing for two hours at 30°C and 220 rpm. The culture was then heat-shocked at 42°C for 15 minutes and incubated for 80 min at 37°C and 220 rpm. Phage were then purified by filtration (0.22 *μ*m). *λ* lysates were stored at −80°C in 15% v/v glycerol. To grow up phage directly from lysates or plaques, ∼2-10 ul of lysate or a picked plaque is added to 4 mL of LBM9 and 100 *μ*L of *E. coli* culture. The culture is then incubated overnight at 37°C and 220 rpm and phage are isolated by centrifugation of host cells at 3900 rcf for 10 min and subsequent filtering of the supernatant (0.22 *μ*m).

### Phage density determination and isolation

Throughout our experiments, we used one of two methods to determine phage densities that both enumerate plaques. Plaques are small clearings in a host bacterial lawn caused the phage population expansion from a single infection. For infused overlays [68], we added 100 *μ*l of a known dilution of phage and 100 *μ*l of a bacterial overnight culture to 4 mL of molten soft agar poured over an LB agar plate. For spot assays [68], 1 mL of bacteria alone was added to 10 mL of molten agar and poured over an LB plate. Once solidified, 2 *μ*l of a known phage dilution was spotted onto the plate. For both methods, plates were incubated overnight at 37°C, plaques were counted, and densities were calculated as plaque forming units per unit volume (pfu/mL). Infused overlays were also used to isolate individual plaques to generated fresh lysates.

### Fluctuating Evolution Experiments

Ancestral *λ* was isolated via infusion overlay plates using LamB^—^ cells. Twelve separate plaques were picked and added to replicate flasks to establish 12 independently-evolving populations. For rounds of evolution towards specialization (Rounds 1 and 3), *λ* was cultured in 10 ml of TrisDM Maltose with 100 *μ*l of OmpF^—^ host (∼1 x 10^9^ cells/mL pre-conditioned in TrisDM Maltose). As the native LamB receptor is a maltoporin, providing maltose as a sugar source leads to the up regulation of LamB. This, in concert with the host lacking OmpF, generates selective pressure to re-specialize on LamB. For rounds of evolution toward generalization (Rounds 2 and 4), *λ* populations were cultured in 10 mL of TrisDM Glucose with 100 *μ*l of wildtype host (∼2 x 10^9^ cells/mL pre-conditioned in TrisDM Glucose). When cultured with glucose, the host down regulates LamB, favoring conditions that select for OmpF function. Wildtype host, rather than L*amB*^—^, was used when selecting for generalization because completely eliminating the native receptor can lead to population extinction. In this way, selection pressure between the two directions was uneven and possibly stronger towards specialization.

For all rounds, flasks were incubated for 24 hours at 37°C at 120 rpm and were serially propagated for 10 days (Rounds 1 and 3) or until all of the populations had regained generalization (5 days for Rounds 2 and 4). Each day, 1 mL from each flask was removed and placed in a 96-deep-well plate. 200 *μ*l of chloroform was added to each well and the plate was incubated at 37°C for 30 min to kill host cells. To initiate the next 24-hour cycle, 100 *μ*l of the previous cycle’s phage stock was transferred to a flask containing fresh media and naive *E. coli* cells growing anew. Each day, a spot assay was conducted with wildtype, LamB^—^, or OmpF^—^ bacteria to assay receptor usage.

At the end of each round of evolution, the chloroformed phage stocks were plated on OmpF^—^ (Rounds 1 and 3 towards specialization) or LamB^—^ (Rounds 2 and 4 towards generalization) to isolate a single plaque representing the endpoint of the experimental round. In this instance, LamB^—^ was used rather than wildtype to ensure the isolated phage remained a generalist. The plaques were grown up overnight with the appropriate host (OmpF^—^ towards specialization, LamB^—^ towards generalization), and the resulting lysates were stored at 4°C and −80°C. 100 *μ*L of this end-point lysate was then used to seed the next round of experiments. In total, 4 rounds of fluctuating evolution were conducted, 2 towards specialization and 2 towards generalization, resulting in 48 strains representing the 12 populations at the end of each of four rounds (Table S6).

### Evolvability Assay

We defined evolvability as the number of days to evolve back to specialization for generalist strains, or back to generalization for specialist strains. To make these measurements, we applied the same experimental conditions used in the fluctuating evolution experiments to the 48 strains, subjecting the 24 strains generated in rounds 2 and 4 to a single round of specializing conditions, and the 24 strains generated in rounds 1 and 3 to a single round of generalizing conditions. We assayed three replicates per strain and evolvability was calculated as the average of the three replicates. We then performed daily spot assays on wildtype, LamB^—^, and OmpF^—^ to monitor receptor usage and calculate evolvability. Strains were considered to have evolved back to specialization once they lost their ability to use the OmpF receptor (e.g., there were no visible plaques present when grown on L*amB*^—^ host) or back to generalization when they regained the ability to use OmpF (*e.g.,* plaques were visible on L*amB*^—^ host).

We conducted the evolvability assays with fresh lysates generated from isolated plaques. For strains generated in rounds 2 and 4 (towards generalization), infused overlays with LamB^—^ host was used to isolate plaques which were subsequently grown overnight with LBM9 and LamB^—^ to generated lysates. Spot plates on wildtype, LamB^—^, and OmpF^—^ were then used to confirm generalization. When plating the lysate from strains from rounds 1 and 3 (towards specialization), we noticed that some strains that had been previously identified as specialists following the fluctuating evolution experiments were now generalists. Given the strains that readily evolve back and forth between specialization and generalization do so rapidly (*e.g.,* in 24 hours), we hypothesized and subsequently confirmed (data not shown) that the rich LBM9 media was causing strains to revert to generalization during plaque amplification. To prevent re-generalization, the freezer stocks from rounds 1 and 3 were first diluted and plated in an infused overlay with OmpF^—^ host. Plaques were resuspended in 12 *μ*l of NaCl, 2 *μ*l was infused in both OmpF^—^ and LamB^—^ to confirm specialization (*e.g.,* plaques only formed on OmpF^—^), and the remaining 8 *μ*l was grown overnight in TrisDM Maltose with OmpF^—^. If the plaques were indeed specialists, the lysate was used to seed the evolvability assays. If the plaques were generalists, the process was repeated 4 more times to confirm those strains were ‘stuck’ as generalists. The strains that remained generalists were excluded from the assay.

For assays in both directions, 100 *μ*l of fresh lysate was added to flasks with 10 mL of TrisDM with the appropriate sugar and host. As for the fluctuating evolution experiments, flasks were incubated for 24 hours at 37°C at 120 rpm and were serially propagated for 10 days (towards specialization) or until all of the populations had evolved back to generalization. Strains were considered to have evolved back to specialization once they lost their ability to use the OmpF receptor (*e.g.,* there were no visible plaques present when grown on LamB^—^ host) or back to generalization when they regained the ability to use OmpF (*e.g.,* plaques were visible on LamB^—^ host). The assays towards specialization and generalization were run separately and were blocked by genotype such that multiple genotypes were assayed simultaneously, but the three replicates were assayed at different times.

### Genomic Sequencing of *λ* Strains

To prepare samples for genome sequencing, fresh lysates were generated from freezer stocks cultured with the appropriate host (OmpF^—^ for Rounds 1 and 3, LamB^—^ for Rounds 2 and 4). For each strain, 500 *μ*L of the lysate was transferred to a 1.5 mL microcentrifuge tube, followed by 50 *μ*L of DNAse Buffer, 10 U of DNAse I (New England Biolabs), and 1 *μ*L of 100 mg/mL RNAse A (Qiagen). The samples were incubated for 30 min at room temperature, at which point 20 *μ*L of 0.5 M EDTA, 2.5 *μ*L Proteinase K (20 mg/mL; New England Biolabs), and 25 *μ*L of 10% SDS were added to each sample. The samples were then mixed and incubated at 55°C for 1 hour. A standard phenol:chloroform extraction was then performed to purify DNA [69]. The aqueous phase resulting from the final chloroform clean-up was transferred to a new tube and an equal volume of 100% ethanol was added. The samples were then added to columns from the PureLink Genomic DNA Kit (Invitrogen Life Technologies) and wash steps were performed according to the manufacturer’s instructions. Water was then used to elute the DNA in the final step. The samples were prepared for sequencing (Illumina NexteraXT), tested for quality, pooled, and sequenced by the UCSD IGM Genomic Center using the Illumina HiSeq4000 to produce 75-bp single-end reads. *breseq* version 0.33.0 was used to align reads to the reference genome and call mutations [70]. For one of the strains, 12R4, coverage was extremely low, as was percentage of mapped reads. We therefore excluded this sample and sequenced the reactive region of J (the informative region of the genome) via Sanger sequencing (see below). For the other strains, coverage of the genome (∼42 kb) was high, with an average coverage of 524, ranging from 243 to 659.

### Sanger sequencing

The reactive region of *J* (host specificity protein, GenBank: NP_040600) was sequenced to confirm genotypes following cloning and genetic engineering and during routine *λ* stock maintenance. J sequencing was also used to generate sequence data for 12R4 and to confirm the presence/absence of stuck mutations and/or reversions in the generated strains. Lastly, we sequenced E. coli *lamB* to confirm the incorporation of the nonsense mutation in our engineered OmpF^—^LamB^—^ double knockout. For these strains, single plaques or colonies were picked and used as template in a standard PCR reaction (Table S7; Q5 High Fidelity 2X Master Mix). Sanger sequencing was performed by Genewiz La Jolla, CA, and samples were submitted as unpurified PCR products and sequenced using the reverse primer. Sequences were aligned using CLC Viewer 7.7.1 and mutations were verified visually.

### Genetic engineering

To test hypotheses regarding the mechanistic basis of the stuck and evolvable mutations, we engineered these mutations alone and in combination into lysogenic phage (c1857) integrated into the *E. coli* genome. This enable us to take advantage of tools developed to direct genomic edits in bacteria (*e.g.,* MAGE). Because the J sequence of cI857 is very different than that of the ancestral *λ*, and these differences could potentially alter the phenotypic behavior of the strain, we first needed to create a cI857 derivative with the ancestral *λ* J sequence. Because λ has an efficient system for homologous recombination [71], we constructed our derivative by first cloning the ancestral *λ* J into a plasmid and then allowing recombination between the plasmid and lysogenic cI857. We first PCR amplified J and ran the product on a 1% agarose gel prepared with SYBR™ Safe DNA Gel stain (Invitrogen). The band was excised and the Wizard® SV Gel and PCR Clean-up System (Promega) to extract and purify the DNA from the gel. We then used the TOPO® TA Cloning® Kit (Invitrogen) to insert the J DNA into the pCR™ 2.1 - TOPO® vector according to the manufacturer’s instructions, using 4 *μ*l of the purified PCR product, 1 *μ*l of the salt solution, and an incubation time of 15 min. We then used Subcloning Efficiency™ DH5*α*™ Competent Cells (Invitrogen) to transform the plasmid according to the manufacturer’s instructions, using 2.5 *μ*l of plasmid vector. Colonies were screened on LB plates with 50 *μ*g/mL ampicillin and 9.5 mg/mL X-gal (5-bromo-4-chloro-3-indolyl-b-D-galactopyranoside; APExBIO) and Sanger sequencing of the J was used to confirm cloning and successful transformation of the ancestral *λ* sequence.

To initiate recombination of the J region, 100 *μ*l of cI857 phage was grown-up overnight with 100 *μ*l of transformed DH5*α* in exponential growth. The resulting lysate was diluted and plated on LamB^—^ to select for transformant phages, as unmodified cI857 is reliant on LamB and unable to produce plaques on this strain. Recombinants were further screened using a PCR assay we developed to screen for the relatively rare recombination in a large number of plaques. We designed a pair of forward and reverse primers flanking the *J* recombination, as well as a forward primer only present within the ancestral recombination (Table S7). After performing a multiplex PCR and visualizing with gel electrophoresis, we could distinguish between phage with the recombination (multiple product) and without (single product). After initial screening, the presence of the recombination was confirmed via Sanger sequencing of the *J*.

Once the ancestral recombination was inserted in the cI857 background, the phage was integrated into the chromosome of *E. coli* strain HWEC106 by inoculating the phage onto a lawn HWEC106 plated on LB agar with 50 *μ*g/mL ampicillin. Plates were incubated at 30°C overnight. Cells that are able to grow within clearing represent those that have λ integrated into their genomes and are immune to reinfection. We therefore picked cells from the center of the plaques and spread them onto LB agar with 50 *μ*g/mL ampicillin and grew at 30°C overnight to isolate individual colonies. Colonies were PCR screened to find a successful lysogen with a single *λ* genome inserted in the canonical *attB* site [72].

We then used MAGE [49], which uses 90-mer single-stranded DNA fragments (see Table S8) to direct genomic edits. We constructed lysogens with the canonical stuck mutation, an evolvable mutation, and both mutations (Table S6). As a MAGE control, we also generated a strain that had gone through the MAGE process, but for which mutations had not been inserted (Table S6, ancestral λ’). We used a modified version of the MAGE coselection protocol to improve efficiency [50]. We coselected with an oligo that generated a premature stop codon in *galK* (Table S8). The oligo targeted to *J* was used at 5 μM, and the *galK* oligo was at 0.05 μM. Each transformation was performed four times and *galK−* cells were selected by plating on deoxy-galactose plates incubated at 37°C. Colonies were picked after 72 h, and Sanger sequencing of *J* was used to confirm transformations.

To test if generalist phage strains could use receptors other than LamB*^—^ and* OmpF^—^, we engineered a double knockout *E. coli* strain. We electroporated the plasmid pKD46 (1.8 kV) from the HWEC106 strain into ∼2 x 10^8^ exponentially growing cells of the OmpF^—^ single knockout (Keio JW0912) washed twice with sterilized water. pkD46 harbors the lambda-red recombineering system and has an ampicillin resistance marker that we used to select for transformed colonies [49]. We then used a standard PCR reaction (Table S7; Q5 High Fidelity 2X Master Mix, New England Biolabs) to amplify DNA from an *E. coli* strain that was previously shown to have a *lamB* nonsense mutation (strain ‘19a’ with a T insertion between positions 610 and 611) [73]. We purified the PCR product using the GenElute Plasmid Miniprep Kit (Sigma Aldrich) and used it as a template to insert the mutation into OmpF^—^ transformed with pkD46. To cause recombination of the nonsense *lamB DNA* with the OmpF^—^genome, we followed the typical MAGE protocol for one round [36, 49] adding the *lamB* double stranded DNA without any additional oligos. The efficiency of transformation with double stranded DNA is low, however, we were able to uncover a successful transformant by exposing the bacterial culture to the lytic base strain, cI26, and isolating a resistant colony. The edit was confirmed by Sanger sequencing.

### Competitive Fitness Assays

To measure fitness, we directly competed our constructed lysogens against a genetically marked ancestral *λ* that can be differentiated when grown on a lawn of *E. coli* cells. When *E. coli* cells with a functional lacZ*α* gene metabolize X-gal, they produce a blue compound. However, cells without this functional gene cannot metabolize X-gal. By inserting a functional lacZ*α* gene into the phage genome, the phage complements the cell’s capacity to metabolize X-gal and generates a blue spot where a plaque forms on a lacZ*α*^—^ *E. coli* (strain DH5*α*). To construct a marked version of our ancestral *λ*, we initiated recombination between *E. coli* (SYP042) with a plasmid encoding the phage recombination (R) gene fused to lacZ*α* (Shao and Want 2008, Want eg al 2003) and our constructed ancestral *λ*. 100 *μ*l of both host and phage were cultured overnight in LBM9 with 50 *μ*g/mL ampicillin at 37°C and 220 rpm overnight. The isolated phage progeny were then plated on a lawn of DH5*α* hosts in the presence of X-gal to screen for plaques for which lacZ*α* had been incorporated into the genome. A single blue plaque was picked, suspended in NaCl (0.85 % w/v), and then a dilution series was plated to purify the phage. Lastly, we constructed a lysogenic version of the marked ancestral strain by plating on *E. coli* HWEC106 and isolating and screening colonies for a single *λ* genome insert as previously described (Table S6, Ancestor λ’_lacZ_).

We ran the competitive fitness experiments using fresh lysates and the same media and host as the fluctuating evolution experiments towards specialization (10 mL TrisDM Maltose, 100 *μ*l OmpF^—^ host preconditioned in TrisDM Maltose). We chose these conditions because the stuck mutation most often evolved in this direction. The assays were blocked such that all phage genotypes were assayed on the same day while the two replicates were run on consecutive days. To initiate a competition, we added approximately ∼1.0 x 10^5^ of the marked ancestral phage and an equal amount of one of the constructed strains to each flask. The mixed cultures were incubated at 37°C at 120 rpm for 24 hrs (the period between transfers in the initial experiments) and phage were sampled and enumerated at the beginning and end of that period. Phage were diluted with TrisDM Maltose and multiple dilutions were plated to ensure individual plaques could be clearly counted. Infusion overly plates were made with DH5a host cells, 9.5 mg/ml X-gal, and 0.02 % IPTG (Isopropyl β-D-1-thiogalactopyranoside; Sigma-Aldrich) and were incubated at 37°C for 24 hrs.

We calculated fitness for the phage using the following equation: [Ln (final/initial densities of the constructed lysogen) – Ln (final/initial densities of the marked ancestral lysogen)] / time (1 day). Note that we used the difference, not the ratio, of the two competitors’ realized growth rates during the competition assay according to Burmeister *et al.,* [38]. This calculation is more reliable when there are large differences in growth rates between competitors [74, 75], which we have previously observed in phage experiments.

### Specialization Assays

For the specialization assays, phage densities on multiple host types were calculated to determine the degree to which phage specializes on the LamB receptor. Spot assays were performed by diluting freshly induced lysates in TrisDM Maltose and spotting on LamB^—^ and OmpF^—^ to calculate pfu/mL. We performed the assay on 3 replicate lysates of each constructed phage and plated each dilution twice for technical replication. We calculated Specialization Index, SI, as follows: SI = (pfu/mL_LamB_ – pfu/mL_OmpF_) / (pfu/mL_LamB_ + pfu/mL_OmpF_). The SI can range from –1 to +1, indicating perfect specialization on OmpF and LamB, respectively, with 0 indicating a symmetric generalist.

### Adsorption-Rate Assays

Adsorption-rate assays were performed on fresh lysates according to Meyer *et al.,* [36] with the following modifications: we diluted the lysates, cultured LamB^—^ and OmpF^—^ cells, and performed the assays using TrisDM Maltose to approximate the conditions of our evolution experiments. We performed the assay on nine biological replicates from each constructed strain. However, because of the time-sensitive nature of the assay, adsorption rates to the two receptors were measured independently on different days.

### Receptor Usage Assay

Freezer stocks of generalist evolvable strains generated in Rounds 2 and 4 of the fluctuating evolution experiments were plated in duplicate on LamB*^—^ and* OmpF*^—^* hosts, as well as on our engineered double knockout (LamB^—^OmpF^—^) host. After 24 hrs, plates were visually inspected for plaques and/or clearings to assay receptor usage.

### Statistical analyses

Statistical analyses were performed in JMP (14.2.0) and R (3.5.2). The distributions of the datasets were first assessed with the Shapiro-Wilk W Goodness-of-fit test. The appropriate statistical test was then chosen based on the distribution of the data and the comparison(s) of interest.

