## Supplementary material for "Viral host-range evolvability changes in response to fluctuating selection": Figure S1

| Cycles | Average number of days |
| --- | --- |
| 0 | 6.42857 |
| 1 | 3 |
| 1 | 5.66667 |
| 1 | 2 |
| 1 | 2.66667 |
| 1 | 5 |
| 1 | 2 |
| 2 | 2 |
| 2 | 3.33333 |
| 2 | 2.66667 |
| 2 | 2 |

**Figure S1. Relationship between the number of cycles of fluctuating selection and the time it takes λ genotypes to re-gain specialization.** Each point represents a unique isolate and the y-value is the average of three evolutionary re-play trials. There is a significant negative relationship (F = 5.6212, P < 0.0418), which indicates that λ evolves to become faster at switching the more rounds it is fluctuated.
