## Supplementary material for "Viral host-range evolvability changes in response to fluctuating selection": Table S8

| **Name** | **Mutations introduced** | **Sequence (5’-3’)** |
| --- | --- | --- |
| MAGE1_RC_stuck | c3227a in *J* | C*C*C*C*GACCCGCTGGCATGTCAACAATACGGGAGAACACCTGTACCtCCTCGTTCGCCGCGCCATCATAAATCACCGCACCGTTCATCAGT |
| MAGE5_RV_Rev | a3319g in *J* | A*G*C*G*CCTGTTTCTTAAACACCATAACCTGCACATCGCTGGCAAACGTATACGGCGGAATTTcTGCCGAATACCGTGTGGACGTAAGCGTG |
| galK_mut45_oligo_galk- | t435a *galK* (premature stop codon) | G*C*T*T*CACTGGAAGTCGCGGTCGGAACCGTATTGCAGCAGCTTTAaCATCTGCCGCTGGACGGCGCACAAATCGCGCTTAACGGTCAGGAA |

**Table S8. Oligos used for genetic engineering of** $\boldsymbol{\lambda}$**.** Asterisks indicate phosphorothioated bond and lowercase letters indicate the bases directing mutations in the oligo.
