## Supplementary material for "Viral host-range evolvability changes in response to fluctuating selection": Table S7

**Table S7. Primer sequences**

| **Name** | **Sequence (5’-3’)** | **Anneal. Temp** | **Use** |
| --- | --- | --- | --- |
| JFor | CGCATCGTTCACCTCTCACT |  | Amplification and sequencing of *J* |
| J500F | GATGACGCCGGACAGCACCACAG |  |  |
| J1100F | TCGGATAAGACGTGGACCTA | 65°C |  |
| J1600F | CCGCTGCGTGAGTATCCGTGAGAA |  |  |
| J2100F | ATAACCGCCACGCCGCATCTT |  |  |
| JRev | CCTGCGGGCGGTTTGTCATTT |  |  |
| J_reco_F | CACCTGACCGCAGAAGTC |  | Identification of the incorporation of ancestral $\lambda$ J recombination into cI857 |
| J_reco_R | CTTTCCGTCCGGTGTCAG | 62°C |  |
| JReShortF2 | TCACGCAGACCGTCAATAA |  |  |
| LamB_F | CTCGGCAACGAAACTCAAATC | 65°C | Amplification of E. coli l*amB* for genetic engineering |
| LamB_R | GCTGATAAACAGAGGACGATGA |  |  |
