## Supplementary material for "Viral host-range evolvability changes in response to fluctuating selection": Table S6

**Table S6.** λ **strains used in study**

| **Name** | **Source** | **Relevant Characteristics** |
| --- | --- | --- |
| cI26 | Donald Court, MD | Strictly lytic base strain |
| Ancestor λ | Evolved | cI26 derivative, OmpF^+^ |
| cI857 | I.-N. Wang, NY | λ for genetic engineering |
| 1-12R1 | Evolved | Generated 1^st^ round of evolution (toward specialization), derived from ancestor λ |
| 1-12R2 | Evolved | Generated 2^nd^ round of evolution (toward generalization), derived from round 1 populations |
| 1-12R3 | Evolved | Generated 3^rd^ round of evolution (toward specialization), derived from round 2 populations |
| 1-12R4 | Evolved | Generated 4^th^ round of evolution t(oward generalization), derived from round 3 population |
| Ancestor λ’ | Engineered | cI857 base strain, ancestral control for engineering process |
| Evolvable | Engineered | cI857 base strain with a3319g mutation in J |
| Stuck | Engineered | cI857 base strain c3227a mutation in J |
| Both | Engineered | cI857 base strain with a3319g and c3227a mutations in J |
| Ancestor λ’_lacZ_ | Engineered | cI857 base strain, lacZ^+^, engineered and marked ancestor |
