## Supplementary material for "Viral host-range evolvability changes in response to fluctuating selection": Table S5

**Table S5. *E. coli* strains used in study.**

| **Name** | **Source** | **Relevant Characteristics** | **Use** |
| --- | --- | --- | --- |
| WT | Keio (BW25113) | λ susceptible | Host for evolution experiments and plating |
| OmpF^—^ | Keio (JW0912) | λ susceptible | Host for evolution experiments, overnight cultures, and plating |
| LamB^—^ | Keio (JW3996) | λ susceptible | Host for overnight cultures and plating |
| LamB^—^OmpF^—^ | Engineered | OmpF^—^ base strain with stop codon inserted in LamB | Host for testing receptor expansion |
| DH5$\alpha$ |  | λ susceptible, lacZα^—^ | Plating host for competitive fitness experiments |
| DH5$\alpha$ | Invitrogen | Competent | Cloning |
| HWEC106 | H. Wang, NY | mutS^—,^ λ-red recombineering plasmid pKD46 | Genetic engineering |
| SYP042 | I.-N. Wang, NY | Contains plasmid with $\lambda$ recombination gene (R) fused to lacZ$\alpha$ | Construction of phage with visual marker |
