## Supplementary material for "Viral host-range evolvability changes in response to fluctuating selection": Table S4

**Table S4. Pairwise Z and p-values for Steel-Dwass test comparing adsorption on LamB and OmpF between constructed strains.** Adsorption on the two receptors was measured independently. Significant p-values ($\alpha$ = 0.05) are in bold.

| **Adsorption on LamB** | | | | **Adsorption on OmpF** | | | |
| --- | --- | --- | --- | --- | --- | --- | --- |
| **Sample 1** | **Sample 2** | **Z** | **p-value** | **Sample 1** | **Sample 2** | **Z** | **p-value** |
| Ancestor | Evolvable | 3.5339 | **0.0012** | Ancestor | Evolvable | 3.5339 | **0.0012** |
| Ancestor | Stuck | 3.3590 | **0.0023** | Ancestor | Stuck | 3.5339 | **0.0012** |
| Evolvable | Stuck | -3.5339 | **0.0012** | Ancestor | Both | -3.5321 | **0.0012** |
