## Supplementary material for "Viral host-range evolvability changes in response to fluctuating selection": Table S3

**Table S3. Pairwise Z and p-values for Steel-Dwass test comparing Specialization Index between constructed strains.** Significant p-values ($\alpha$ = 0.05) are in bold.

| **Sample 1** | **Sample 2** | **Z** | **p-value** |
| --- | --- | --- | --- |
| Ancestor | Evolvable | 2.8022 | **0.0261** |
| Ancestor | Stuck | 0.4003 | 0.9783 |
| Ancestor | Both | 2.8022 | **0.0261** |
| Evolvable | Stuck | 2.8022 | **0.0261** |
| Evolvable | Both | 0.5605 | 0.9437 |
| Stuck | Both | -2.8022 | **0.0261** |
