## Supplementary material for "Viral host-range evolvability changes in response to fluctuating selection": Table S2

**Table S2. Pairwise p-values for Tukey-Kramer HSD test comparing competitive fitness between constructed strains.** Significant p-values ($\alpha$ = 0.05) are in bold.

| **Sample 1** | **Sample 2** | **p-value** |
| --- | --- | --- |
| Ancestor | Evolvable | **0.0031** |
| Ancestor | Stuck | **0.0013** |
| Ancestor | Both | **0.0005** |
| Evolvable | Stuck | 0.2514 |
| Evolvable | Both | **0.0171** |
| Stuck | Both | 0.0925 |
